# Segregation of Familial Risk of Obesity in NHANES Cohort Supports a Major Role for Large Genetic Effects in the Current Obesity Epidemic

**DOI:** 10.1101/749606

**Authors:** Arthur B. Jenkins, Marijka Batterham, Lesley V. Campbell

## Abstract

The continuing increase in many countries in adult body mass index (BMI kg/m^2^) and its dispersion is contributed to by interaction between genetic susceptibilities and an increasingly obesogenic environment (OE). The determinants of OE-susceptibility are unresolved, due to uncertainty around relevant genetic and environmental architecture. We aimed to test the multi-modal distributional predictions of a Mendelian genetic architecture based on collectively common, but individually rare, large-effect variants and their ability to account for current trends in a large population-based sample. We studied publicly available adult BMI data (n = 9102) from 3 cycles of NHANES (1999, 2005, 2013). A first degree family history of diabetes served as a binary marker (FH_0_/FH_1_) of genetic obesity susceptibility. We tested for multi-modal BMI distributions non-parametrically using kernel-smoothing and conditional quantile regression (CQR), obtained parametric fits to a Mendelian model in FH_1_, and estimated FH x OE interactions in CQR models and ANCOVA models incorporating secular time. Non-parametric distributional analyses were consistent with multi-modality and fits to a Mendelian model in FH_1_ reliably identified 3 modes. Mode separation accounted for ~40% of BMI variance in FH_1_ providing a lower bound for the contribution of large effects. CQR identified strong FH x OE interactions and FH_1_ accounted for ~60% of the secular trends in BMI and its SD in ANCOVA models. Multimodality in the FH effect is inconsistent with a predominantly polygenic, small effect architecture and we conclude that large genetic effects interacting with OE provide a better quantitative explanation for current trends in BMI.

## Introduction

The recent and continuing increase in the global mean adult BMI, first seen in high income countries, is now seen in most countries across a wide range of ethnic composition and socio-economic conditions (Di Cesare et al., 2016) and is accompanied by increases in measures of dispersion (Krishna et al., 2015; Silventoinen et al., 2017). Although BMI is known from family-based studies to be under strong genetic influences (Loos, 2018) population genetic backgrounds have been effectively constant over this time, implying that BMI trends are driven by change in environmental factors (obesogenic environment, OE). Evidence from twin studies, which demonstrate increased genetic variance over time, supports an important role for interactions between OE and genetic susceptibility (G x OE) on both mean and dispersion of BMI (Rokholm et al., 2011; Silventoinen et al., 2017), but how large a role is not yet known. Defining the role of G x OE in “epidemic” obesity, and hence of genetic susceptibility itself, is hindered by problems of measurement and modeling of interactions (Franks and McCarthy, 2016) and by uncertainty around both the genetic architecture (Loos, 2018) and the exact nature of the environmental drivers (Hall, 2018). Whether population susceptibility to OE is predominantly determined by a subgroup with high genetic susceptibility or is more evenly spread within populations is unresolved despite important implications for the management of obesity and related disorders at population and individual levels (Kivimaki et al., 2015; Krishna et al., 2015; Jenkins and Campbell, 2015; Razak et al., 2015).

The genetic variants responsible for obesity susceptibility remain largely unknown. Genome-wide association studies (GWAS) have identified significant associations with >200 markers with small effects on BMI (polygenes), together explaining only approximately 3-4 % of total variance compared to family-based heritabilities (h^2^) of 50-75% (Speakman et al., 2018). Few causative mechanisms responsible for these phenotypically weak associations are known (Loos, 2018). The sources of the h^2^ unaccounted for by GWAS are uncertain; suggestions include overestimation of h^2^, large numbers of common genetic variants with small, statistically insignificant effects on phenotypes (Locke et al., 2015; Khera et al., 2019) and importantly, candidates not tested in most GWAS. Among the latter are rare genetic variants with large phenotypic effects and G x OE interactions (Loos, 2018). Recently significant G x OE interactions have been detected in individual GWAS loci and in composite genetic risk scores, which however explain little of the missing component of h^2^ (Abadi et al., 2017; Nagpal et al., 2018).

A family history of diabetes (FH) is a potent, predominantly genetic (Hemminki et al., 2010; Willemsen et al., 2015) risk factor for diabetes diagnosis (DM) and for obesity-related phenotypes (Ghosh et al., 2010; Tirosh et al., 2011; Scott and Consortium, 2013; Jenkins et al., 2013) consistent with the strong association between type 2 DM and overweight/obesity. Familial effects on obesity-related phenotypes in adults are also predominantly genetic (Stryjecki et al., 2018; Silventoinen et al., 2017), so to the extent that the DM generating FH is of type 2 (approximately 94% of DM in the US population (Xu et al., 2018)), FH is a prevalent and readily obtained marker of genetic susceptibility both to diabetes and to the obesity commonly preceding it. We have previously reported evidence from a small sample of a multi-modal effect of FH on a composite adiposity index consistent with segregation in families of discrete obesity risk (Jenkins et al., 2013). Polygenic risk scores (PRS) based on large numbers of small effects are expected to be, and appear to be, unimodally-distributed (Llewellyn et al., 2014; Rask-Andersen et al., 2017) and thus cannot account for familial segregation of discrete risk. The present work is based on the alternative hypothesis that individually rare, but collectively common, genetic variants with large phenotypic effects are the source of most of the missing h^2^ and of most of G x OE, and that their effects can be detected through analyses of phenotypic segregation in high-risk families (Jenkins and Campbell, 2014).

The Continuous National Health and Nutrition Examination Survey (NHANES) is a continuing (1999-) large-scale population-based survey incorporating an index of adiposity (Body Mass Index, BMI) and first-degree FH (FH_0_/FH_1_) together with potential covariates and confounders. Although BMI has recognized limitations as an adiposity phenotype (Jenkins and Campbell, 2014; Müller et al., 2018) it is the basis for most large-scale genetic studies and like other authors, we assume that a large enough scale and appropriate modeling of covariates will reduce effects of imprecision and bias (Speakman et al., 2018). We aimed to test in a large multi-cycle NHANES sample for the presence of familial segregation of genetic risk and to estimate the contribution of FH, and by extension all discrete genetic risk, to recent secular trends in adult BMI. The results support a predominant role for large genetic effects interacting with OE in the obesity “epidemic”.

## Subjects and Methods

### Subjects

We used data from the 1999-2000, 2005-2006 and 2013-2014 cycles of NHANES (https://www.cdc.gov/nchs/nhanes/index.htm accessed 25 Aug 2017). We extracted records for participants age 20-65 years with non-missing gender, race/ethnicity, BMI, diabetes family history (FH) and diabetes diagnosis data, and current smoking status if available. The definitions of two fields changed over the sampling period: 1) Diabetes family history was defined in terms of 1° and 2° relatives in 1999-2000 but by 1° relatives only in subsequent cycles. We recoded the 1999-2000 diabetes family history data to conform to the later definition using the separately collected data for affected parents and siblings. 2) The self-identified race/ethnicity field (RIDRETH1) code was used excluding other races (OR) to maintain consistency of categories across cycles (Supplementary Methods). We excluded from the primary analyses subjects diagnosed with diabetes because of possible confounding by effects of either diabetes or diabetes therapies on BMI. The resulting data set is summarized in Table 1.

**Table 1:** Participants by diabetes status (DM_0_/DM_1_)

|  | NHANES cycle |  |  |  |  |
| --- | --- | --- | --- | --- | --- |
|  | All cycles | 1999-2000 | 2005-2006 | 2013-2014 | p<sup>†</sup> |
| **DM<sub>0</sub>** |  |  |  |  |  |
| n | 9102 | 2865 | 3076 | 3161 | - |
| Gender (F%) | 53 | 55 | 54 | 52 | 0.06 |
| Race/Ethnicity (%)<br>(MA/OH/NHW/NHB)<sup>‡</sup> | 23/8/46/23 | 30/7/44/20 | 24/4/49/24 | 16/11/48/25 | 7.8 x 10<sup>-56</sup> |
| Age (yr): Mean | 40.5 | 40.7 | 39.4 | 41.5 | 4.5 x 10<sup>-9</sup> |
| SD | 13.2 | 13.2 | 13 | 13.2 | 0.63 |
| BMI (kg/m<sup>2</sup>)*: Mean | 28.7 | 28.3 | 28.6 | 29.2 | 4.6 x 10<sup>-9</sup> |
| SD | 6.6 | 6.3 | 6.4 | 7.1 | 1.2 x 10<sup>-11</sup> |
| Diabetes Family History<br>(Y%) | 36 | 29 | 42 | 38 | 4.1 x 10<sup>-25</sup> |
| Current smoking<br>(%, Y/N/missing) | 25/20/55 |  |  |  | - |
| **DM<sub>1</sub>** |  |  |  |  |  |
| n (%) | 793 (8.0) | 211 (6.9) | 252 (7.6) | 330 (9.5) | 0.0003 |
| Age at Diagnosis (yr) | 43.6 | 44.3 | 42.9 | 43.6 | 0.47 |

† Cycle effects (p) by ANOVA (age), ANCOVA (BMI), Bartlett's test (SD's) and Chi-squared test for categorical
variables.
‡ MA = Mexican American, OH = Other Hispanic, NHW = Non-Hispanic White, NHB = Non-Hispanic Black
\* Adjusted for age, gender and race/ethnicity in a linear model (see Table S1).

### Statistical analyses

#### Approach

We treated the data as a convenience sample and took no account of the sampling weights provided by NHANES to permit nationally representative estimates. Our results are not intended to be representative of the US population.

Our primary analyses are based on non-parametric visualization (kernel-smoothing) and analyses (conditional quantile regression, CQR) of distributions requiring no prior distributional assumptions. Parametric fits to multimodal distributions were then used to quantify the contributions of the predicted large genetic effects model. FH(_0/1_) is treated analytically as a binary genetic risk marker but the distribution of its effect across quantiles is interpreted in a Mendelian model in which FH_1_ represents enrichment of a mixture of single and double carriers of risk variants. Calendar time is treated as a continuous surrogate of OE. Effects of OE interacting with FH were assessed in CQR models, and also in least-squares ANCOVA models using bootstrap resampling to minimize distributional assumptions in the calculation of effect size estimates and errors. All analyses were performed using R 3.6.1 (R Development Core Team, 2016).

#### Summary statistics

Heterogeneity of the samples across cycles was assessed by Chi square test for categorical variables and by one-way ANOVA for age. Effects on BMI were assessed by ANCOVA against continuous time (yr = calendar start year - 1999). Effects on phenotype SD’s were assessed by Bartlett’s test in one-way ANOVA models.

#### Adjustment

Prior to analysis BMI was adjusted for effects of age, gender and race/ethnicity in a linear model (age + age^2^ + race x gender). The adjustment model accounted for 4 % of the total variance in BMI (Supplementary Table S1).

#### Distributions

##### Visualization

The effect of FH on the distribution of adjusted BMI was visualized using kernel-smoothed density estimates by FH status (R base function density). The degree of smoothing is controlled by the bandwidth parameter (bw) which was obtained in the full non-diabetic data set (bw = 0.99) from a measure of the dispersion of the data (Sheather and Jones, 1991). This produces a continuous distribution function and is widely used to visualise features of potential interest which may be obscured in histograms. The credibility of the apparent effect of FH on the shape of the distribution was assessed by post-hoc analysis of density ratios (FH_1_ / FH_0_) by quantiles (20) of the full sample. Mean density ratios with SEM were obtained by quantile by bootstrap resampling with replacement (1000 draws, stratified by FH status with resample sizes = strata sizes) and compared to the predictions of normal and log-normal mixture distributions characterised by the proportions, means and SD’s in the sample stratified by FH status and cycle. Lack of fit to the mixture distributions was assessed by X^2^ tests in 1/variance-weighted linear regressions.

##### Conditional Quantile Regression (CQR)

Conditional quantile regression is a powerful tool for analyzing the effect of covariates on distributions without assumptions of distributional shape. In contrast to ordinary least-squares (OLS) regression which characterizes effects on global features of a distribution, CQR analyses local effects of covariates independently at any specified quantiles and can detect variations in covariate effects across quantiles. Originating in econometrics (Koenker, 2017) it is now used in other areas including genetics (Briollais and Durrieu, 2014). In the CQR framework developed by Abadi et al (Abadi et al., 2017) for analysis of genomic markers, trends in effect sizes across quantiles represent interactions between genetic effects and unobserved environmental and/or genetic factors. We treat FH as a binary genetic risk marker (FH_0_/FH_1_) and a linear trend in quantile regression coefficients (ß_1i_) across quantiles (**τ**_i_) represents summed linear interactions of FH with unobserved factors. We analysed the effects of FH on adjusted BMI by CQR using the R package quantreg. The effect of all interactions on the FH effect was tested in in a 2 parameter linear model:
for each quantile **τ**_i_ in y,

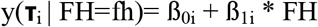

where y(**τ**_i_ | FH=fh) = the ith quantile of adjusted BMI conditional on the value of FH (0,1), the intercept ß_0i_ is the ith quantile value in FH_0_ and ß_1i_ is the FH effect size in quantile i.

The interaction between FH and continuous calendar time was estimated in the ANCOVA model:

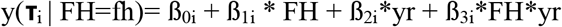

where yr = cycle start year – 1999, ß_0i_ is the ith quantile value in FH_0_ at yr =0 and ß_2i_ and ß_3i_ are CQR coefficients for time and time*FH interaction effects in quantile i. The coefficients for the time-related effects represent the effects in FH_0_ (ß_2i_) and the additional effects in FH_1_ (ß_3i_) so that the coefficients for total time effects in FH_1_ = ß_2i_ + ß_3i_.

Equality of CQR parameter effect sizes across quantiles was tested using the anova.rq function in the R package quantreg.

The strengths of the CQR effects across quantiles were also assessed in linear meta-regression analyses of relationships between quantile coefficients ß_1-3i_ and quantiles using the R package metafor (Abadi et al., 2017). Regression coefficients (ß_MR_) with SEM are reported in units of kg.m^−2^ over the full quantile scale (0-1). The structure of the FH effects in relation to the multi-modal Mendelian hypothesis was assessed in an analysis of residuals from linear OLS fits of ß_1i_ to ß_0i_ with Durbin-Watson tests for residual 1^st^ order autocorrelation (function durbinWatsonTest in the R package car).

##### Parametric fits

We obtained fits to a 3-component normal mixture distribution representing a simple Mendelian model of fixed genetic effect using an expectation-maximization algorithm (normalmixEM function in the R package mixtools). The models are characterised by the fitted means (μ_i_), standard deviations (σ_i_) and mixing proportions (λ_i_) of the three component distributions. Full model fits were obtained in FH_1_ but were not obtainable in FH_0_ or DM_1_ groups and we constrained μ_i_ in those groups to values estimated in FH_1_ in order to obtain comparable estimates of σ_i_ and λ_i_. Risk allele frequencies (q) under an additive Mendelian model of large genetic effects were calculated from the fitted λ_i_:

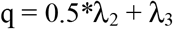

where λ_2_ and λ_3_ represent the proportions of carriers of 1 and 2 risk alleles respectively. Within-sample consistency of calculated q across the three groups analysed was assessed by comparing fitted q_FH_1__ with the prediction from random mating of DM_1_ into the full sample:

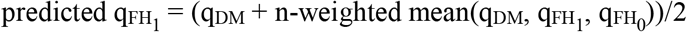

#### Secular trends

Effects of diabetes family history status (FH_0_, FH_1_) and continuous time (yr = calendar start year - 1999) on adjusted BMI and its standard deviation (SD) were assessed in ANCOVA models of the form:

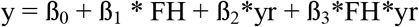

where y = adjusted BMI mean or SD by FH status (0/1) and cycle, and yr = cycle start year – 1999. Each OLS fit estimated 4 parameters from the 6 data points. Mean parameter estimates with 95% CI were obtained by bootstrap resampling with replacement (1000 draws stratified by FH status and cycle with resample sizes = strata sizes).

#### Comparison of cross-sectional and secular trend effects

Effects of FH on BMI distribution and on secular trends in BMI were compared by calculating the contribution (%) of FH_1_ to the effect in the full non-diabetic sample for calculated risk allele frequency (q%) and to the slope (ß%) of the relationships between time and BMI in ANCOVA model described above. Mean (± SEM where possible) q% and ß% were calculated in the relevant bootstrap samples.

## Results

### Participant characteristics

Data from 9102 non-diabetic subjects met the inclusion criteria, approximately equally distributed across the 3 cycles. Gender balance varied little but there was a cycle effect in race/ethnicity, most obvious in the reduced representation of MA in the two later cycles. Average age varied across cycles but not its SD, while adjusted BMI and its SD showed linear trends with cycle time. FH_1_ prevalence was higher in the two later cycles compared to 1999-2000 as was DM_1_ prevalence. Current smoking status was predominantly missing in the data (55%) and was not included in the BMI adjustment model. However smoking status was not related to FH status whether analysed in the full data (X^2^ = 2.80, 2 df, p = 0.25) or in those with non-missing smoking status (X^2^ = 0.43, 1 df, p = 0.51), hence is unlikely to confound analyses of FH effects. The mean age at diagnosis of DM (43.6 yr) is consistent with predominant type 2 DM in the sample.

### Distributions

#### Visualization

Adjusted BMI in the non-diabetic sample showed an apparently unimodal distribution, right-skewed compared to a normal model and closer to a log-normal model (Fig 1A). When visualized by FH status (Fig 1B) the predicted multimodality in FH_1_ was indicated with modes in the normal weight, overweight and obese regions of the BMI distribution. In contrast FH_0_ showed an apparently unimodal distribution. A difference in shape between the two groups was supported by the analysis of density ratios (Fig 1B inset) in which models based on mixtures (FH x cycle) of either normal or log-normal distributions did not provide adequate fits to the data (p ≤ 0.001). BMI distribution in the diabetic sample appeared to be depleted in the lower mode and enriched in the upper modes compared to FH_1_ (Fig 1C).

**Figure 1:**
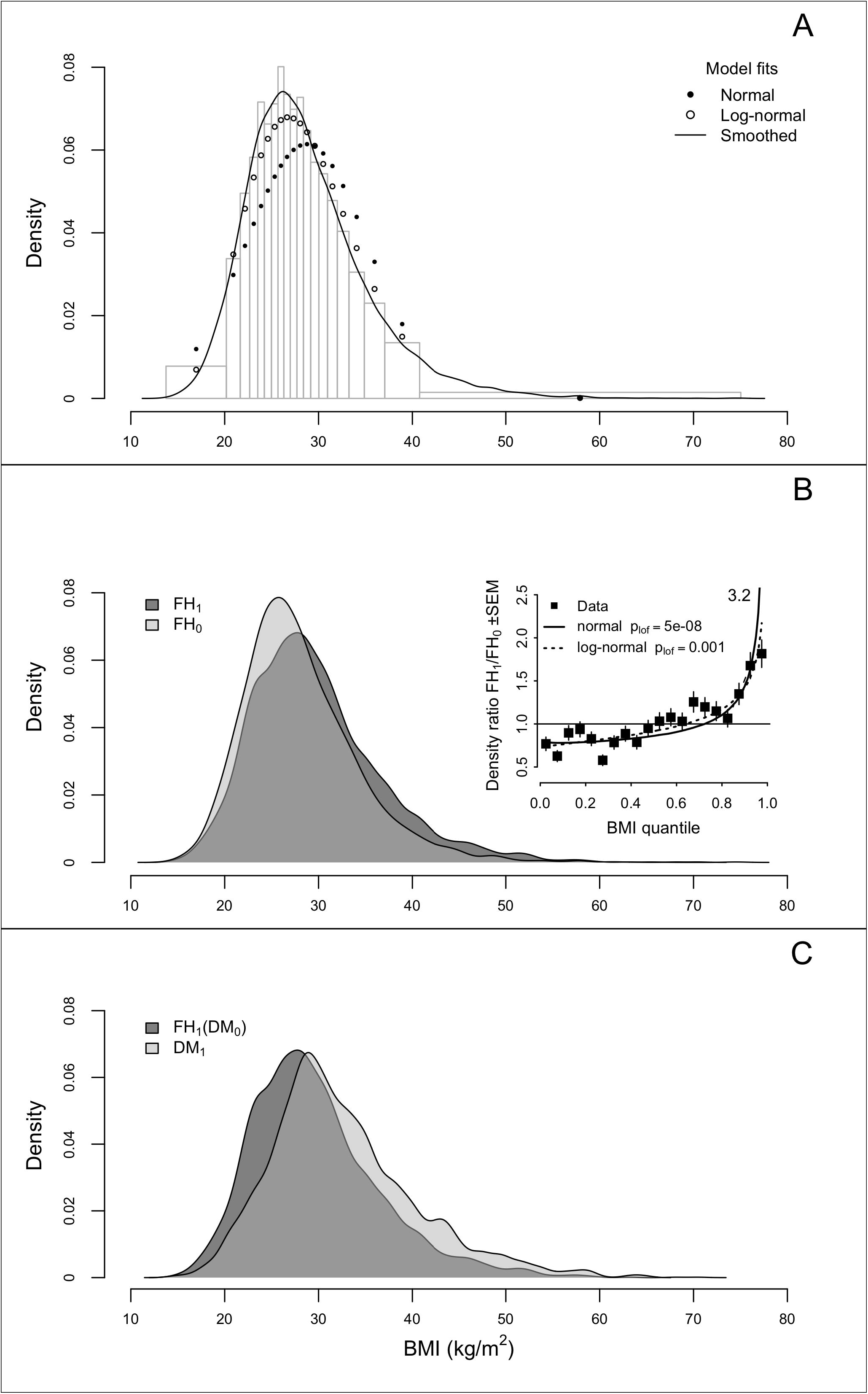
Distribution of adjusted BMI in non-diabetic (DM_0_) and diabetic (DM_1_) participants in combined NHANES 1999-2000, 2005-2006 and 2013-2014 cycles. A) Full non-diabetic sample (n=9102) binned by quantiles (n=20) with superimposed kernel-smoothed and fitted densities in normal and log-normal models. B) Main: kernel-smoothed adjusted BMI density by FH status. Inset: density ratios (FH_1_ / FH_0_) ± SEM obtained by bootstrap resampling by quantiles of the full sample compared to predictions of normal (solid line) and log-normal (dotted line) mixture models with p associated with lack-of-fit testing (p_lof_). C) Kernel -smoothed adjusted BMI density in non-diabetic FH_1_ (n=3297) compared to diabetic participants (n=793).

#### CQR

Analysis of the effect of FH status alone on the shape of the BMI distribution using CQR demonstrated increasing FH effect size at higher levels of BMI (ß_1MR_ = 2.2 ± 0.2 (SEM) kg.m^−2^ Fig 2A, main panel), indicating strong interactions between FH_1_ and other variables not included in the model. FH_1_ effect size ranged from < 1 kg/m^2^ in the lower quantiles to ~3 kg/m^2^ in the upper quantiles, substantially different in both regions to the OLS estimate (1.7 kg/m^2^). Inclusion of calendar time in the two-way model weakened the trend in ß1 across quantiles (ß_1MR_ = 1.5 ± 0.3, Fig 2B main panel) and exposed significant OLS effects and trends across quantiles for main (ß_2MR_ = 0.07 ± 0.02) and interaction (ß_3MR_ = 0.11 ± 0.03) effects of time (Fig 2C-D, main panels). The total OLS and MR interaction effect sizes in FH_1_ (ß_2OLS_ + ß_3OLS_ = 0.11 ± 0.03 (SE), ß_2MR_ + ß_3MR_ = 0.18 ± 0.04) were approximately double those in FH_0_ (ß_2OLS_, ß_2MR_, Fig 2C). The analysis supports the conclusion that calendar time is a strong surrogate of OE interacting with genetic factors as represented by FH status, and that susceptibility to OE increases with increasing BMI in both groups but more strongly in FH_1_.

**Figure 2:**
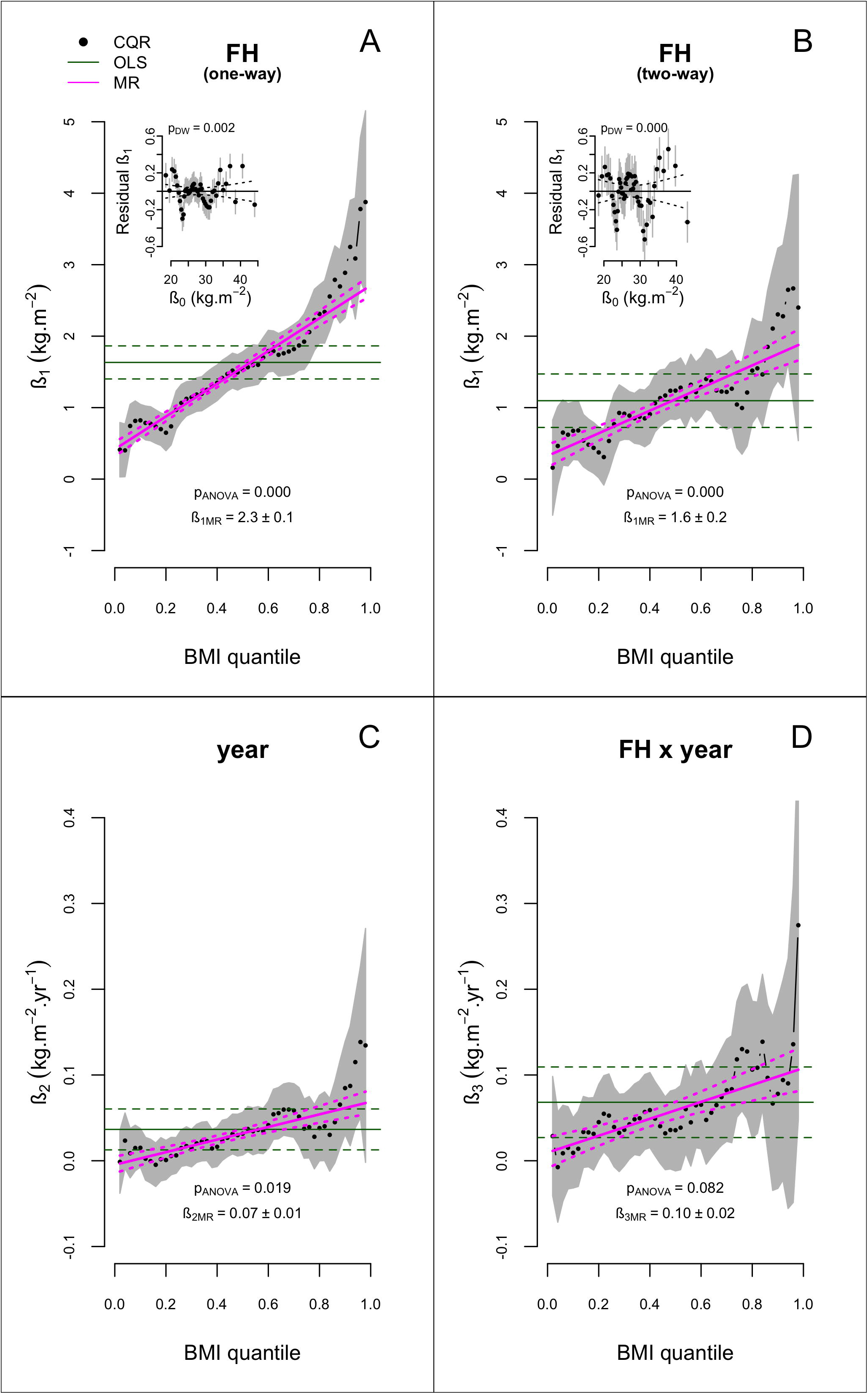
Conditional quantile regression effects of diabetes family history (FH) on adjusted BMI in non-diabetic participants in models consisting of FH alone (A) and in interaction with continuous calendar time (B-D). The main panels show the effect sizes (ß_1-3_) by quantile with 95% CI (grey-shaded area), the OLS estimate of the average effect (solid green line) with 95% CI (dotted green lines) with p-values (p_ANOVA_) from anova tests for equality of ß_i_ across quantiles and the meta-regression fits ± 95%CI (magenta lines) with ß_MR_ ± SEM. The insets in panels A and B show the patterns of residuals (Δß_1i_) ± residual SEM from linear OLS fits of ß_1i_ (FH_1_) to ß_0i_ (FH_0_) with 95% CI on the fits (dotted lines) around the lines Δß_1_ = 0 representing perfect fits, and with the p value (p_DW_) from a Durbin-Watson test of autocorrelation of residuals. CQR estimates of the SEM of ß_0i_ are also depicted but are mostly obscured by the point symbols.

While the overall MR relationships between ß_1_ and quantiles in both models were approximately linear (Fig 2A,B) there was strong evidence for additional non-linear structure in the OLS analysis of ß_1i_ against ß_0i_ (insets Fig 2A,B). The linear models provided good fits (one-way: slope = 0.15 ± 0.003 (SE), R^2^ = 0.98; two-way: slope = 0.10 ± 0.01, R^2^ = 0.86) but residual sequential structure was apparent in both models, confirmed by tests of autocorrelation in residuals (p_DW_ ≤ 0.002). The pattern of residuals in the two-way model (Fig 2B inset), adjusted for time-related effects, shows clear signs of discrete effects of FH_1_ on the distribution of adjusted BMI with distinct peaks in the lower, middle and upper regions of the distribution. This pattern in the conditional quantile coefficients does not map directly onto the unconditional quantile plots in Fig 1B, but does highlight similar regions in the distribution, and permits the conclusion that FH_1_ has discrete, not continuous effects on BMI.

#### Parametric analysis

The distribution of BMI in FH_1_ (Fig 1B) and the discrete pattern in the FH_1_ effect by CQR (Fig 2B inset) appear consistent with a simple Mendelian model and fitting a 3-component normal distribution model to the FH_1_ data resulted in robust estimates of component means (Fig 3A) and mixing coefficients and SDs (Table 2). Approximately 50% of FH_1_ occupied the upper two modes and separation between modes accounted for approximately 40% of the total variance in adjusted BMI with the remainder assigned to dispersion within modes (Fig 2B). Under a Mendelian model the variance due to mode separation represents a lower bound on the contribution of large effects as some of the dispersion within modes represents variance in effect sizes of individual contributing causal loci (see Discussion) which will contribute to the ~60% of variance assigned to within-modes. Estimates of component SDs and mixing proportions with component means, constrained for FH_0_ and DM_1_ to those identified in the FH_1_ data (Table 2), support enrichment in the two upper components in FH_1_ compared to FH_0_ (48% vs. 33%) and more strongly in DM_1_ (72%). Predicted risk allele frequencies in FH_1_ (q - Table 2) express these distributional properties in Mendelian terms and show within-sample consistency in that q_FH1_ predicted from random mating of DM_1_ (0.37) is not different to the fitted estimate (0.30 ± 10).

**Table 2:** Three-component normal mixture distribution fits to adjusted BMI by FH and DM status^¥^.

|  |  | DM<sub>0</sub> |  | DM<sub>1</sub> |
| --- | --- | --- | --- | --- |
|  | Component | FH<sub>0</sub> | FH<sub>1</sub> |  |
| mean | 1 | * | 25.8±1.0 | * |
|  | 2 | * | 32.1±1.8 | * |
|  | 3 | * | 40.6±2.5 | * |
| sd | 1 | 3.8 | 3.8±0.9 | 3.7 |
|  | 2 | 5.7 | 5.1±1.2 | 4.5 |
|  | 3 | 10 | 8.2±0.8 | 8.4 |
| λ | 1 | 0.67 | 0.52±0.16 | 0.28 |
|  | 2 | 0.29 | 0.37±0.15 | 0.47 |
|  | 3 | 0.04 | 0.11±0.06 | 0.25 |
| q<sup>†</sup> | - | 0.18 | 0.30±0.10 | 0.49 |
|  |  |  | (0.37)<sup>§</sup> |  |

<sup>¥</sup> ± bootstrap standard error for FH<sub>1</sub> only524 \* Means in FH<sub>0</sub> and DM<sub>1</sub> constrained to fitted values in FH<sub>1</sub>525 <sup>†</sup> Calculated risk allele frequency in an additive Mendelian model of large effects526 <sup>§</sup> Predicted from DM<sub>1</sub> mating randomly into the full sample.

**Figure 3:**
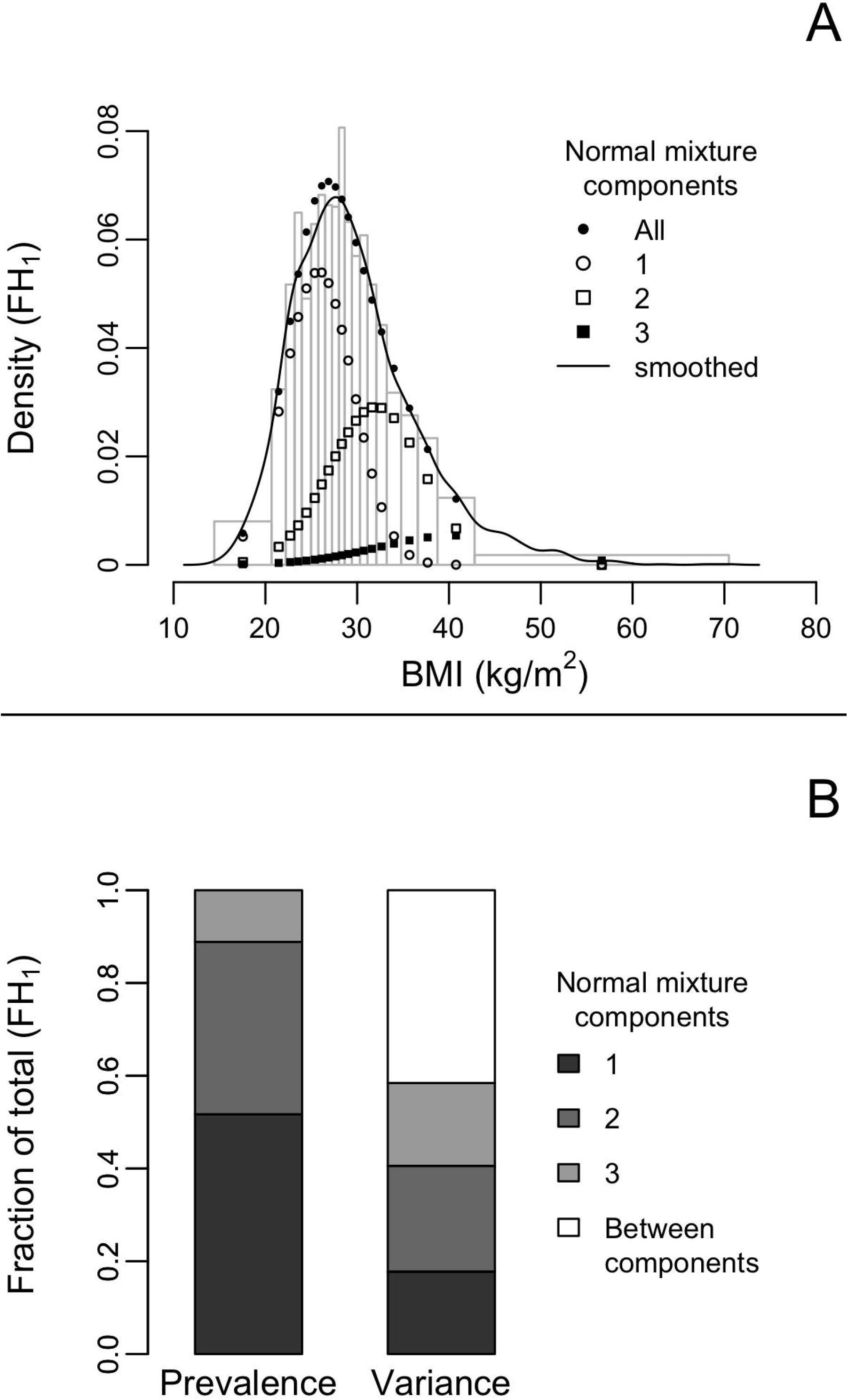
A) Adjusted BMI density in FH_1_ by quantile (grey bars) and kernel-smoothed (black line) with fits to a three component normal mixture distribution. B) Estimated contributions in FH_1_ of the components of the mixture distribution to the prevalence (mixture coefficients, λ) and variance of adjusted BMI.

**Figure 4:**
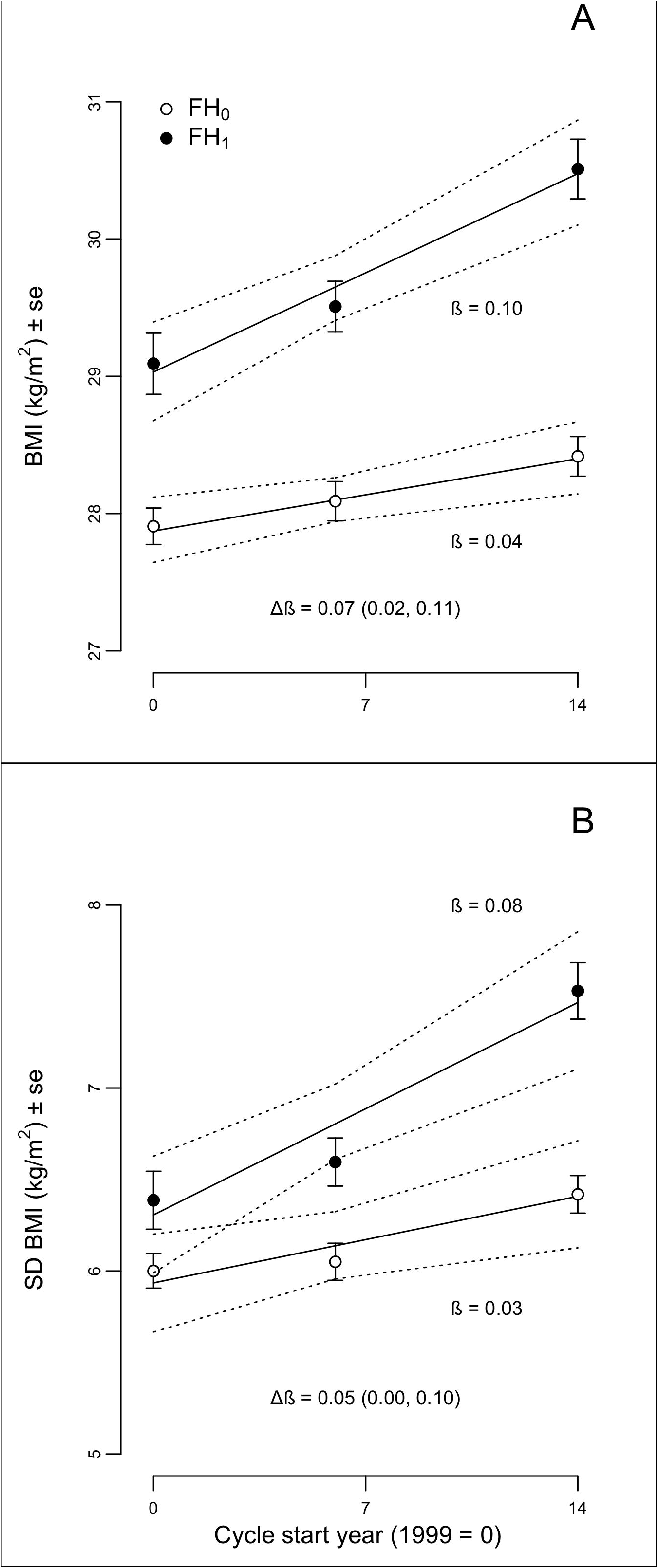
Effects of diabetes family history (FH_0/1_) on linear secular trends in age-, gender- and race/ethnicity-adjusted BMI mean ± sem (A) and standard deviation ± sem (B). Parameter estimates with 95% CI were obtained in ANCOVA models by stratified bootstrap resampling of all non-diabetic individuals (see Methods). Dotted lines enclose 95% CI on fitted values at each point; ß = regression slope vs. time (kg/m^2^ per year); Δß = ß_FH1_ − ß_FH0_ (95%CI).

### Secular trends

Adjusted BMI mean (Fig 2A) and SD (Fig 2B) increased over the sampling period significantly faster in FH_1_ compared to FH_0_ in the bootstrapped ANCOVA model, and estimates of ß and Δß in the mean data were indistinguishable from the OLS estimates provided by the CQR analysis (Fig 2C,D). Similar results were obtained with log-transformed BMI (Supplementary Fig S2). FH_1_ accounted for 62% of the BMI mean trend and 60% of the BMI SD trend in this sample over the period 1999-2014, effects similar in magnitude to the estimated FH_1_ contribution to the sample risk allele frequency (50%, Supplementary Table S2).

## Discussion

### Summary

We tested the prediction of segregation of discrete effects of FH on adult BMI (Jenkins et al., 2013), modeled as modes of distribution, and estimated the contribution of FH_1_ to recent trends in BMI mean and dispersion in a large population-based sample. The results support a predominant role in the recent obesity “epidemic” for rare genetic variants with large effects interacting with OE.

### Segregation of genetic susceptibility

The non-parametric analysis provided evidence for a multi-modal distribution in the FH_1_ group consistent with the prediction of segregation of large genetic effects in families (Jenkins et al., 2013). Multi-modality was supported by the analysis of density differences between FH_1_ and FH_0_ by unconditional quantiles (Fig 1B) and by evidence of discrete signals in the OLS analysis of CQR coefficients (Fig 2A&B insets). The two upper peaks in Fig 2B inset are consistent with the predicted Mendelian effects of large effect variants on BMI but the potential lower peak is unexpected and may reflect the presence of type 1 diabetes family history in FH_1_ group. Individuals with type 1 diabetes often present with BMI in the underweight (<18.5) – normal range (<25) (Manyanga et al., 2016) but a genetic basis for this has not been established.

Polygenic risk scores (PRS) in population-based samples are expected to be unimodally-distributed, and appear to be so (Llewellyn et al., 2014; Rask-Andersen et al., 2017). Any elevated polygenic obesity risk in DM_1_ will dilute into the mating population resulting in a right-shifted distribution in FH_1_ compared to FH_0_, not discrete effects. Alternative explanations for multi-modality might be discrete stratification of OE components not captured by calendar time which seems unlikely, or un-modeled interactions between FH and other covariates. Un-modeled interactions between FH and stratified residual confounders may exist and contribute but we found no evidence for this in plots of distributions by gender and race/ethnicity (Supplementary Fig S1) or in an analysis of smoking status against FH. Discrete inheritance of genetic variants with large effects is the most likely explanation for multi-modality in the FH effects on the BMI distribution.

Approximately 40% of the adjusted BMI variance in FH_1_ was accounted for by between-modes variance (Fig 3B) but this represents a lower bound since the identified modes are likely to be synthetic ie composed of a range of effect sizes due to rare variants at different loci. Indications of fine structure within the broad central peak (Fig 2B inset) are suggestive. Examples of rare variants with large effects on BMI in adults (ß) are known from studies of candidate genes and monogenic obesity loci (Jenkins and Campbell, 2014) while more recently a common variant in Samoans (EAF = 0.26, ß ≈ 1.4 kg/m^2^), very rare in other populations (Minster et al., 2016), and an African-specific rare variant (EAF = 0.008, ß =4.6 kg/m^2^) undetected in Europeans and Asians (Chen et al., 2017) have been identified by GWAS. Overall, ß in these nine examples ranges from 1.4- 9 kg/m^2^ and a similar range in effect sizes in the NHANES sample would contribute substantially to the within-mode variance estimated here. A combination of within-subject variance (~5% (Wormser et al., 2013)) with polygenic variance (~ 5% (Loos, 2018)) sets a lower bound for within-modes variance and hence the upper bound for between modes, implying that between 40% and 90% of total variance in FH_1_ may be attributed to large genetic effects.

### G x OE

FH_1_ is a prevalent (36%) and powerful determinant of the rate of change of mean BMI and its dispersion over time in the ANCOVA models, accounting for 62% of the BMI trend and 60% of the BMI SD trend in this sample over 1999-2014. Consistent results were obtained in the CQR models with ß_OLS_ and ß_MR_ in FH_1_ approximately double those in FH_0_. Under a polygenic model the familial risk would be distributed normally over FH_1_ which would then be a marker of a large fraction of the at-risk population. However under the Mendelian model supported here genetic risk would segregate in families and only approximately 50% of FH_1_ would acquire the excess familial risk and only ~18% of the sample would then account for ~60% of the trends. Individuals with a family history DM_1_ must represent a fraction of individuals with elevated genetic obesity risk and it is likely that the remainder, particularly those with a family history of obesity without DM, would substantially increase the genetic component of the trends consequent to the high heritability of BMI (Stryjecki et al., 2018). This Mendelian model is internally consistent in estimates of risk allele frequencies (q) in FH_1_, FH_0_ and DM_1_ (Table 2) and in comparisons of FH_1_ effect sizes in cross section (q, ~50%) and over time (ß, ~60%) (Supplementary Table S2). Our results support the proposition that the largest part, and perhaps all, recent trends in mean and dispersion of BMI are due to a minor subset of individuals with elevated genetic susceptibility to OE.

### Limitations

The design and interpretation of fits to parametric mixture distribution models involves choices concerning the number of components, parameter starting values and algorithms, and fit to a specific model cannot be taken in isolation as support for its structural validity. We base our choice and structural interpretation of 3-component normal mixture model fits and parameters on the *a priori* hypothesis of Mendelian segregation of obesity risk in families (Jenkins et al., 2013) supported by the non-parametric distributional plots (Fig 1B,C) and CQR analysis (Fig 2A,B insets). Like Abadi et al (Abadi et al., 2017), we interpret interactions in the CQR analysis as predominantly G x OE although a contribution from G x G interactions cannot be excluded. Our interpretation is supported by the effects on the interaction of including calendar time in the CQR model (Fig 2B). Other limitations discussed above include our inability to exclude discreet stratification of OE and the possible influence of unmeasured/unknown confounders of FH effects.

### Conclusions

We conclude that a Mendelian model of individually rare but collectively common genetic risk variants with large effects interacting with OE provides a plausible quantitative explanation for recent trends in obesity and should be favored over a predominantly polygenic model which does not. The evidence for a predominant role for polygenes (Khera et al., 2019) can appear to be strong (eg “Polygenic obesity is the most common form of obesity in modern societies…” (Albuquerque et al., 2017)) but recent interpretations seek to explain the still missing heritability in obesity in terms of unidentified large genetic effects and G x OE (Saeed et al., 2018; Loos, 2018) and recommend a renewed focus on family-based designs and on specific populations in which large effect variants may be enriched and identified(Minster et al., 2016; Chen et al., 2017). Our results strengthen that view by showing that a model based on unidentified segregating variants with large effects interacting with OE can account for the largest part of the secular trend in BMI and its dispersion in a large population-based sample.

## Author Contributions Statement

All authors contributed to the study design. AJ extracted and analysed the data in consultation with MB. AJ and LC wrote the first draft of the manuscript. All authors contributed to manuscript revision and editing.

