## Supplemental data for "Segregation of Familial Risk of Obesity in NHANES Cohort Supports a Major Role for Large Genetic Effects in the Current Obesity Epidemic" (from another version of this preprint: version 2, 2019-11-15)

### Supplementary Methods

All cycles included the self-identified race/ethnicity field (RIDRETH1) but the 2013 cycle also included a redefined field (RIDRETH3) in which those who identified as non-Hispanic Asian (NHA) received a separate coding, having in previous cycles been coded as other race (OR). As NHA were oversampled by design in 2013 and were coded OR in the 2013 RIDRETH1 field the composition of the OR group in RIDRETH1 changed in 2013 compared to the earlier cycles. To maintain consistency of definitions across cycles we therefore used the RIDRETH1 race/ethnicity data after exclusion of OR.

**Supplementary Table S1: BMI-adjustment model parameters in combined NHANES cycle data (n = 9102, $R^2$ = 0.04)**

| Parameter | | Estimate ($kg/m^2$) ± sem | p |
|---|---|---|---|
| age | | 59.6 ± 6.5 | **<0.001** |
| $age^2$ | | -43.8 ± 6.5 | **<0.001** |
| Male | | | |
| | MA* | 28.42 ± 0.21 | **<0.001** |
| | OH | -0.36 ± 0.43 | 0.402 |
| | NHW | -0.40 ± 0.25 | 0.121 |
| | NHB | -0.30 ± 0.30 | 0.305 |
| Female | | | |
| | MA* | 0.98 ± 0.29 | **0.001** |
| | OH | -0.39 ± 0.58 | 0.502 |
| | NHW | -0.84 ± 0.35 | **0.016** |
| | NHB | 2.34 ± 0.40 | **<0.001** |

* Reference categories within and between genders: Female MA estimate = MA(Female-Male)

**Supplementary Table S2: Comparison of effects of diabetes family history ($FH_0$ – n = 5805 , $FH_1$ – n =3297) on secular trends in adjusted BMI and its SD and on calculated risk allele frequencies (q) in non-diabetic participants.**

| Measure | FH status | Estimate ± SE | %Total[¥] ± SE[*] |
|---|---|---|---|
| BMI trend* | $FH_1$ | 0.10 ± 0.02 | 62 ± 11 |
| | $FH_0$ | 0.04 ± 0.01 | |
| BMI SD trend* | $FH_1$ | 0.08 ± 0.02 | 60 ± 15 |
| | $FH_0$ | 0.03 ± 0.02 | |
| q[†] | $FH_1$ | 0.30±0.10 | 50 |
| | $FH_0$ | 0.18 | |

* Linear rates of change in BMI and BMI SD ($kg.m^{-2}.yr^{-1}$) in $FH_1$ and $FH_0$ across cycle years ± standard errors (SE) calculated by bootstrap resampling in the analysis summarized in Fig 2A.

[¥] group size weighted $FH_1$ estimate/ Total (%) with Total calculated as group size weighted sum of $FH_0$ and $FH_1$ estimates.

[†] Risk allele frequency calculated under an additive Mendelian model from fits to 3-component normal mixture distributions summarised in Table 2. Comparable SE not obtainable in unconstrained fits for $FH_0$ and hence in %Total.

### Supplementary Fig S1: Kernel-smoothed adjusted BMI densities by FH, gender and race/ethnicity

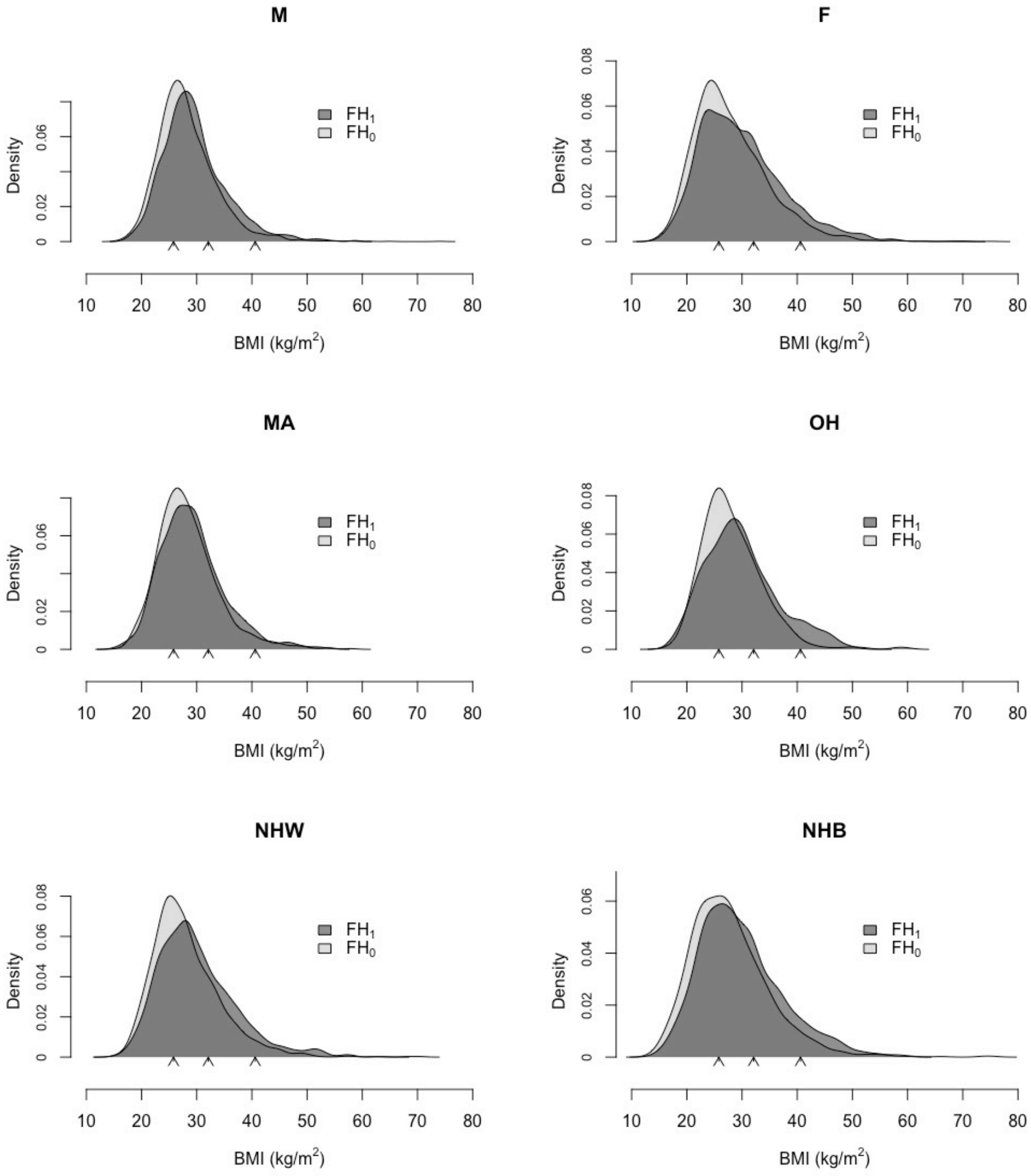


Kernel-smoothed adjusted BMI density by FH status (n/group) within genders (M, F) and race/ethnicity categories (MA, OH, NHW, NHB). Red arrows mark the fitted component means in the 3-component normal distribution fits to the full FH+ data summarized in Table 2.

### Supplementary Fig S2: secular trends in log-transformed adjusted BMI

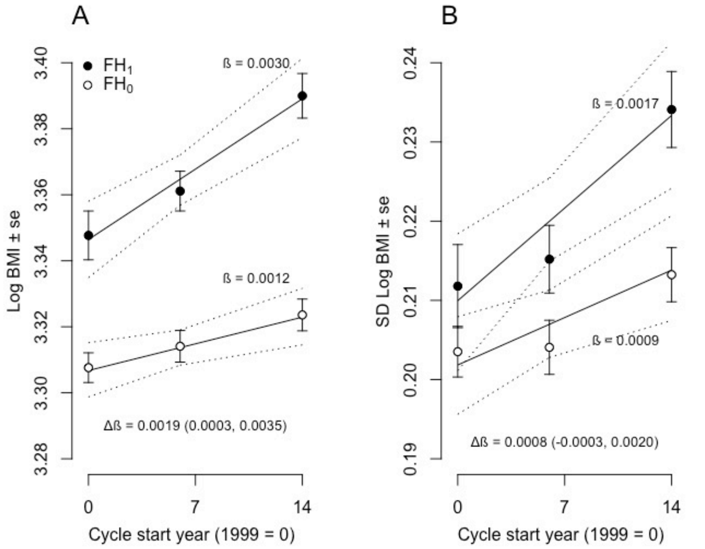


Effects of diabetes family history ($FH_{0/1}$) on linear secular trends in log-transformed age-, gender- and race/ethnicity-adjusted BMI mean ± se (A) and standard deviation ± se (B). Estimates with 95% CI were obtained in ANCOVA models by stratified bootstrap resampling of all non-diabetic individuals (see Methods). Dotted lines enclose 95% CI on fitted values at each point; ß = regression slope vs time (kg/m$^2$ per year); $\Delta ß = ß_{FH1} - ß_{FH0}$ (95%CI).
